# Data-driven spectroscopic dictionaries and detector-calibrated inference for photon-limited Raman hyperspectral imaging of living cells

**DOI:** 10.64898/2026.08.31.748229

**Authors:** Shunsuke Yagi, Norihide Sagami, Ikuto Eshima, Kotaro Hiramatsu

## Abstract

Label-free Raman imaging of living cells is photon limited: at exposures compatible with cellular dynamics, single-pixel spectra carry about one count per channel on a dominant smooth background. We present an unmixing framework in which the decoder of a physics-constrained autoencoder is restricted to a data-driven spectroscopic dictionary: band centers, widths, and pseudo-Voigt shapes are measured from the dataset and fixed, and the network learns only nonnegative band amplitudes, a smooth B-spline background, and a per-pixel gain. First, on slit-scanning images of HeLa cells (532 nm) the dictionary yields spike-free component spectra that read as band tables, including a resonance-enhanced cytochrome-*c*-associated component matching literature spectra, and the most stable decomposition against the component number. Second, the dictionary and initialization calibrated at 1 s exposure per line transfer to 100 ms per line (*~* 12 s sweeps): cytochrome-*c* spectral identity survives a single sweep (correlation 0.92) while its map remains photon limited; the dictionary provides spectral physicality, and the transferred initialization prevents a structural collapse that global map correlations miss; in a measurement-derived phantom the dictionary estimator holds the cytochrome-*c* spectrum to 17–19° spectral angle at 100 ms, where classical factorizations and free decoders lose it (55–64°). Estimation on the count-equivalent detector output uses a calibrated shifted-Poisson quasi-likelihood. Third, evaluation must be time matched: correlation against a separately acquired reference saturates through slow specimen drift and acquisition mismatch rather than photon noise, and the self-consistency of learned denoisers is inflated by shared bias; time-matched self-consistency and independent cross-checks are proposed.

## I. INTRODUCTION

Spontaneous Raman microscopy provides label-free chemical contrast in living cells, but the Raman cross-section is small: even with slit-scanning excitation^1,2^ and resonance enhancement of selected chromophores^3^, the photon budget at exposures short enough to follow cellular dynamics is of order one detected count per pixel and spectral channel. Under these conditions the two standard steps of hyperspectral analysis, background removal and spectral unmixing, become strongly coupled and ill-posed. The continuum background (medium fluorescence plus a broad cellular pedestal) carries most of the detected energy; the chemically informative bands are small structures on top of it; and the per-pixel noise is Poisson-dominated with a non-trivial detector transfer for analog electron-multiplying CCDs (EMCCDs)^4^.

Classical matrix-factorization approaches, such as multivariate curve resolution (MCR-ALS)^5^, nonnegative matrix factorization (NMF)^6^, and pure-pixel extraction of endmembers (the component spectra) by vertex component analysis (VCA)^7^ or N-FINDR^8^, can be applied after denoising and baseline correction^9,10^, but at low photon counts the background dominates every factor, pure-pixel methods select noise outliers, and the factorization becomes rotationally ambiguous. Autoencoder (AE) unmixing^11^ improves scalability and allows physical constraints in the decoder, yet a decoder that can synthesize arbitrary nonnegative spectra absorbs baseline into “chemical” components or, when regularized toward positive peaks on a regular grid, produces artificial combs of narrow spikes at positions where the data contain no band at all.

The central idea of this work is to separate what should be *measured* from what should be *learned*. The positions, widths, and line shapes of the Raman bands present in a dataset are measurable spectroscopic quantities: we extract them once, from representative spectra of the dataset itself, by a bounded multi-peak pseudo-Voigt fit^12^ seeded by asymmetrically-reweighted baseline estimation^9^. The resulting *data-driven dictionary* is then frozen inside the decoder of a chemical-plus-background autoencoder, and the network learns only (i) nonnegative band amplitudes for each chemical component, (ii) per-pixel abundances on a simplex with a positive intensity gain, and (iii) a smooth, low-dimensional background field. Known physics (illumination gain fields, detector calibration) is applied analytically in the forward model; unknown noise is handled by a quasi-likelihood whose parameters are calibrated on the data and whose limits we state.

We develop and test this framework on slit-scanning Raman images of HeLa cells at 1 s and 100 ms exposure and of C2C12 myoblasts at 1 s, all at 532 nm excitation, within the Q-band absorption of reduced cytochrome *c*, where its heme modes are resonance enhanced^3,13^. Beyond the decomposition method itself, the low-photon, live-cell setting forced us to re-examine how such methods should be *evaluated*. Two failure modes of standard practice turned out to be quantitatively important: (i) correlation against a reference image from a separate acquisition is limited by specimen drift and acquisition mismatch, not by the estimator, so that apparent “irrecoverability” of a component can be an artifact of the evaluation; and (ii) the self-consistency of learned denoisers such as Noise2Noise^14^ is inflated by shared model bias and must be audited against independent estimators that do not share the learned bias. Both effects are demonstrated explicitly, and simple protocols (time-matched interleaved self-consistency; partial-correlation crosstalk audits; band-resolved fidelity metrics) are proposed.

## II. EXPERIMENTAL DATA AND DETECTOR MODEL

### A. Datasets

Line-illumination (slit-scanning) hyperspectral Raman images were acquired at 532 nm excitation with an imaging spectrograph equipped with an EMCCD detector (Andor iXon DU897; details below). Each camera frame is a 512× 512 (*y, λ*) image of the illumination line; scanning the line in *x* (0.25 µm steps) builds the hyperspectral cube. Three datasets are used: (i) HeLa, 1 s exposure per line, one sweep of 100 positions; (ii) HeLa, 100 ms exposure, ten consecutive sweeps (time-lapse) of the same field; (iii) C2C12 myoblasts, 1 s exposure, four fields of view. The wavenumber axis was calibrated on polystyrene beads; the usable window is 530–1750 cm^−1^ (448 channels) after discarding the illumination edge. The along-line sampling is 0.0968 µm per row. The first 15 scan positions of every sweep contain a shutter transient and are discarded, and cosmic rays are replaced using a temporal median (time-lapse data) or a spectral median filter (single sweeps).

Because the illumination line is scanned, an image is not a snapshot. The 1 s image is a rolling acquisition of 102 s (kinetic cycle 1.022 s per position: 1 s exposure plus 21 ms readout); a 100 ms sweep takes 12.2 s (cycle 121.8 ms); and the ten-sweep time-lapse spans 122 s. According to the acquisition timestamps in the file headers, the 1 s image of the HeLa field was recorded first and the 100 ms time-lapse was started 7.6 min later, i.e. about 6 min after the end of the 1 s acquisition. These time scales are essential for interpreting map comparisons across acquisitions (Sec. IV D).

#### Instrument and sample details

The 532 nm excitation line (300 mW at the sample) was focused and the Raman scattering collected through a 100 ×oil-immersion plan-achromat objective (PLN100XO, Evident; NA 1.25, working distance 0.15 mm). The spectrum of each point along the line was dispersed by an imaging spectrograph (MK-300, Bunkoukeiki; F/4.4) with a 600 grooves/mm grating, giving 2.72 cm^−1^ per detector column, onto the EMCCD (Andor iXon DU897, 512 ×512 pixels, frame-transfer mode, EM-gain setting 300 (a DAC value, not the gain), kinetic cycle equal to the exposure plus 21 ms readout). HeLa cells (RIKEN BRC Cell Bank, RCB0007) were cultured in MEM with 10% calf serum and C2C12 myoblasts (RIKEN BRC, RCB0987) in low-glucose DMEM with 10% fetal bovine serum, both at 37 °C under 5% CO_2_ following the supplier’s protocols.

### B. Detector calibration and noise model

The EMCCD is operated in analog EM mode, so the output is not photon-counting. We use the count-equivalent scale *W* = (*D −B*)*/k* with bias *B* = 150 raw units (from signal-free pixels), read noise *σ*_*r*_ = 16.2 raw units, and *k* = 105.1 raw units per count obtained by *second-moment matching* of the variance–mean relation of the data, var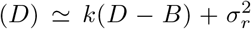 (statistical uncertainty *±*0.2; systematic range 98–111 depending on the pixel population used). The scale *k* absorbs the EM excess-noise factor (*k ≃ F*^2^ ≃ *g* with *F*^2^≃1.86, *g* ≃ 56.5^4^). We do not assign a probability law to the continuous output *y*. Instead we specify only its first two moments, the mean and the *variance function*,

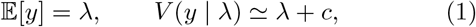

where *c* = (*σ*_*r*_*/k*)^2^ = 0.0238 in count-equivalent units^15^, i.e. the variance function of a Poisson variable shifted by *c*.

A two-moment specification of this kind is precisely what Wedderburn’s *quasi-likelihood* ^16,17^ requires: the objective is built from the variance function rather than from a density. The quantity to be fitted is the mean *λ*, so the natural estimating function is the mean residual *y− λ* divided by its own variance, and the objective is its antiderivative in *λ*,

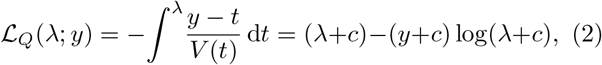

summed over pixels and channels (*λ*-independent constants of integration omitted). Equation (2) has the algebraic form of the negative log-likelihood of a Poisson variable shifted by *c*—hence “shifted-Poisson”—but no such law is asserted for *y*; only Eq. (1) is.

What this objective does to a fit is seen most directly in its derivative. For a forward model *λ*_*i*_(*θ*) with parameters *θ* (in Sec. III B, the network weights, the dictionary amplitudes, and the gain fields entering *λ*_*i*_), differentiating Eq. (2) by the chain rule gives

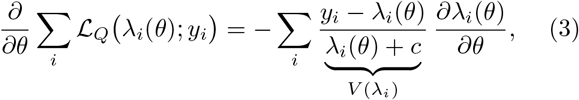

so that at the optimum Σ_*i*_[(*y*_*i*_−λ_*i*_)*/V* (*λ*_*i*_)]*∂*_*θ*_*λ*_*i*_ = 0, Wedderburn’s quasi-score equation. Each residual therefore enters weighted by the reciprocal of its own variance, evaluated at the current fitted mean: on these data a bright background channel (*λ*_*i*_ ≃30 counts) is down-weighted by a factor *~*90 relative to a faint band channel (*λ*_*i*_ *≃*0.3 counts). Ordinary least squares is the special case *V ≡* const, which weights every residual equally regardless of intensity. The weights are not free parameters and are not estimated from repeats: they are fixed by the calibrated variance function and follow the fit. The correspondingly standardized residuals are the Pearson residuals 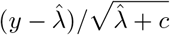.

The construction also delimits what is assumed. The minimizer of Eq. (2) is consistent whenever the mean model *λ*_*i*_(*θ*) is correct, whatever the true law of *y*; misspecification of moments beyond the second costs efficiency and invalidates nominal standard errors, but not consistency^16,17^. That is what licenses Eq. (2) on an EMCCD output that is manifestly not Poisson, provided Eq. (1) holds to second order—which is the property the repeat measurements are used to test.

Two properties of Eq. (1) are fixed by the calibration above and are therefore not tests of it: the unit slope of the variance–mean relation and, to first order, the unit standard deviation of the Pearson residuals of the quasi-likelihood fits described below (measured 1.02; the same residuals under a homoscedastic Gaussian model have standard deviation 1.41, which quantifies the heteroscedasticity that a least-squares fit ignores). What the data do test is (i) linearity of the variance in the mean over the full intensity range with a single scale *k*, which holds on the 100 ms repeats, and (ii) the third cumulant of the repeat distribution, which the second-moment calibration leaves free. Measured on the ten 100 ms repeats (binned on an independent intensity estimate to avoid selection bias), the third cumulant scales as *κ*_3_ ≃ (1.5–1.7) *λ* over 0.3–30 counts, i.e. it follows the electron-multiplication cascade (*κ*_3_ = 6*λ/F*^4^ = 1.5– 1.7 *λ* for *F*^2^ = 2–1.86^4^) rather than the Poisson value *κ*_3_ = *λ* or the Gaussian value zero; the variance floor is 0.09 counts^2^, larger than the nominal read-noise term *c* but exceeding 10% of the variance only below*~* 0.9 counts per channel (Appendix B, Fig. 9). These measurements thus support Eq. (1) as a second-moment specification while excluding the corresponding probability law, which is exactly the regime in which the quasi-likelihood argument above applies; we refer to the minimizers of Eq. (2) as quasi-likelihood (not maximum-likelihood) estimators throughout. The nominal *c* rather than the measured floor is used in the objective because *c* affects only the weighting of the faintest channels: refitting the raw-domain quasi-likelihood estimator of Sec. IV B with *c* = 0.09 (or *c* = 0) instead of 0.024 changes the component map correlations by at most 0.03 after resolution matching, with no consistent sign, and brings the held-out Pearson residual standard deviation from 1.02 to 1.00; the measured floor is thus preferable for residual calibration but immaterial for the estimates. The modest Poisson-versus-least-squares differences reported in Sec. IV B are consistent with this status. The overall workflow built on this detector model is summarized in Fig. 1.

**FIG. 1.**
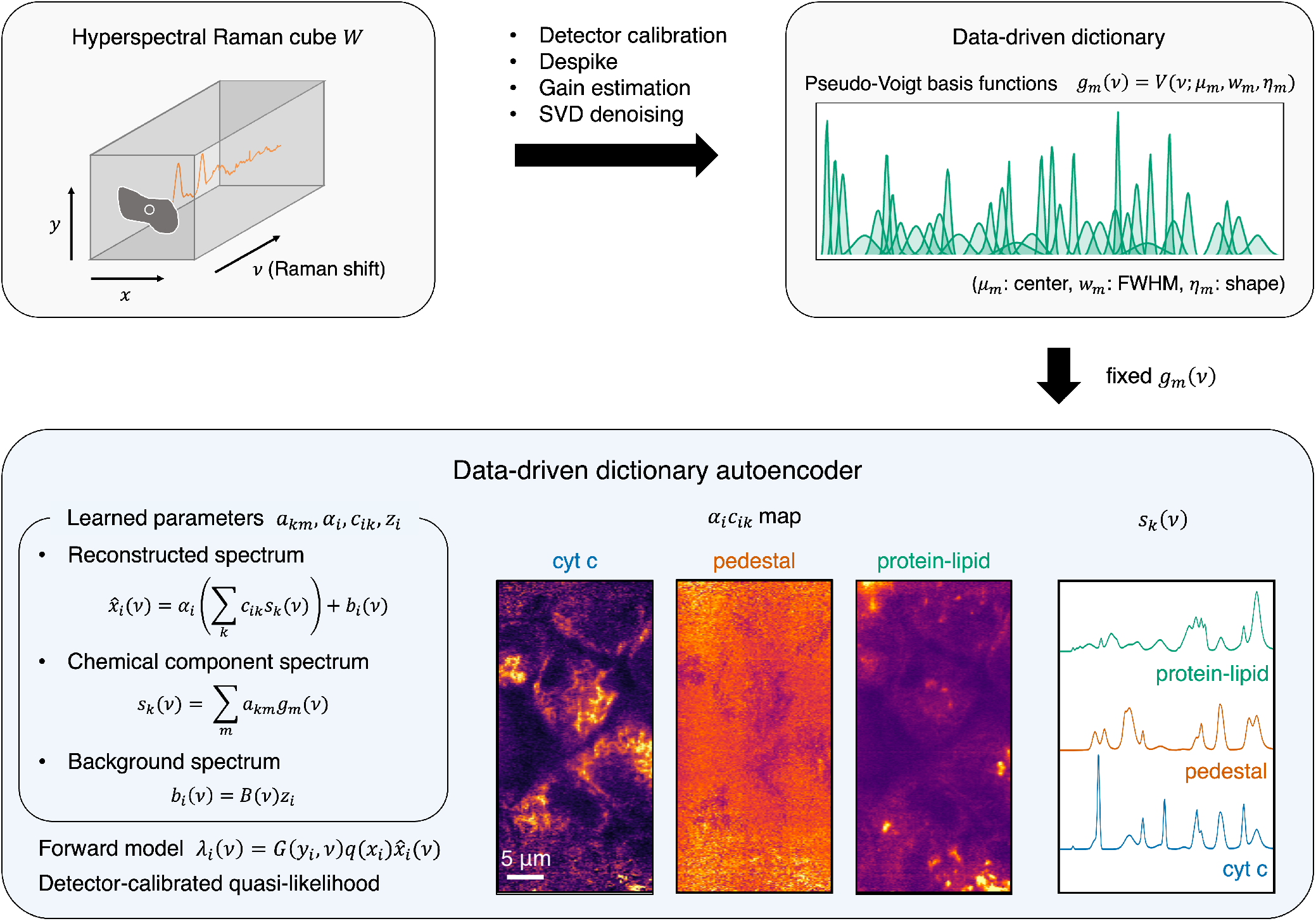
Analysis workflow. The measured hyperspectral cube (*x, y, ν*), rendered from the 1 s HeLa data as the integrated intensity followed by baseline-subtracted band images at 751 cm^−1^ (cytochrome *c*) and 1660 cm^−1^ (amide I) and an off-band continuum channel (each voxel column holds one noisy single-pixel spectrum, inset), is despiked and converted to count-equivalent units, and the illumination gains *G*(*y, ν*), *q*(*x*) are estimated from cell-free continuum. A data-driven dictionary (band centers, widths, and pseudo-Voigt shapes measured from representative spectra of the dataset itself) is frozen inside the decoder of the chemical-plus-background autoencoder, Eq. (5), which learns only band amplitudes *a*_*km*_, simplex abundances, a per-pixel gain, and a smooth B-spline background. Quasi-likelihood estimators operate on the undivided count-equivalent output with the variance function of Eq. (1) and the gain fields in the forward model. The output is a set of chemical amplitude maps (cytochrome *c*, pedestal, protein–lipid; scale bar 5 µm).

### C. Illumination-gain correction

Two multiplicative instrument fields are estimated from cell-free continuum and treated as known in all subsequent analysis. They are estimated once per acquisition series from the data with the highest signal-to-noise ratio (SNR) available (the sweep average for the 100 ms time-lapse): a single low-dose sweep is too photon-starved for a reliable mask-free estimate, and applying gains estimated from it corrupts the data. The two fields are: a row-by-wavenumber gain *G*(*y, ν*) (line-illumination speckle, the smooth illumination envelope along the line, and three low-sensitivity detector rows, recovered at their expected 8–25% depth) and a static scan-direction profile *q*(*x*) (*±*9%). Both are estimated mask-free from a low percentile across scan positions. Analysis is restricted to the well-illuminated band of rows (envelope *>* 0.25 of maximum), because dividing by a small gain would otherwise amplify line-edge noise into the statistics. When a quasi-likelihood estimator on the undivided count-equivalent output is used the data are *not* divided by these gains; instead the gains multiply the forward model, preserving the count statistics. (Dictionary construction, an auxiliary measurement of band positions and widths, is performed on the gain-corrected representation denoised by singular-value decomposition (SVD); see Sec. III A.)

## III. METHODS

### A. Data-driven dictionary construction

Representative spectra are formed from the (gain-corrected, SVD-denoised) cube: the mean spectrum, the per-channel 95th percentile, and the per-channel standard deviation. The percentile and dispersion spectra are essential: several bands of spatially localized species are invisible in the global mean (seven of the bands of the final HeLa dictionary, including the 928 cm^−1^ pedestal band and the tyrosine doublet, were detected only in the percentile or standard-deviation representatives).

For each representative, an airPLS baseline^9^ provides an *initial* background estimate, and peaks are detected in the residual with prominence, distance, and width thresholds specified in cm^−1^. Candidates from all representatives are merged (| Δ*ν*| *<* 5 cm^−1^, prominence-weighted centers). The merged set is then refined by a bounded multi-peak fit to the 95th-percentile spectrum,

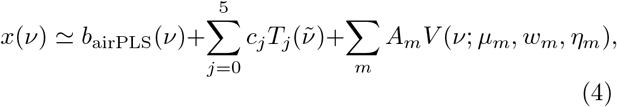

where *V* is an area-normalized pseudo-Voigt^12^ with center bounds 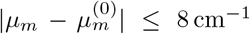, full width at half maximum (FWHM) *w*_*m*_ ∈ [5, 120] cm^−1^, *η*_*m*_ ∈ [0, 1], and *A*_*m*_ ≥0. The Chebyshev term is a *bounded* local readjustment of the airPLS baseline (total correction*≤* 15% of the baseline maximum); without the bound the baseline and the broadest peaks exchange roles, an instability we observed directly. After the fit, the residual is rescanned for missed candidates (this recovered the weak 604/624/644 cm^−1^ bands and resolved the 1311/1318 cm^−1^ doublet), weak peaks are pruned by relative area, and the final dictionary *{µ*_*m*_, *w*_*m*_, *η*_*m*_*}* is frozen. For the HeLa 1 s data this yields *M* = 38 bands (*R*^2^ = 0.9996 for the representative fit) with directly recognizable assignments: the cytochrome-*c* resonance fingerprint (751, 1130, 1317, 1586 cm^−1^), phenylalanine (1005), CH deformation (1455), amide I (1660), amide III (1239/1258), the tyrosine doublet (823/857), a nucleic-acid band (787), and one broad (FWHM *≃*61 cm^−1^) cellular pedestal band at 928 cm^−1^ that is deliberately retained on the chemical side [Fig. 2]. The widest bands of the dictionary reach FWHM *≃*92 cm^−1^ (1070 and 1390 cm^−1^).

**FIG. 2.**
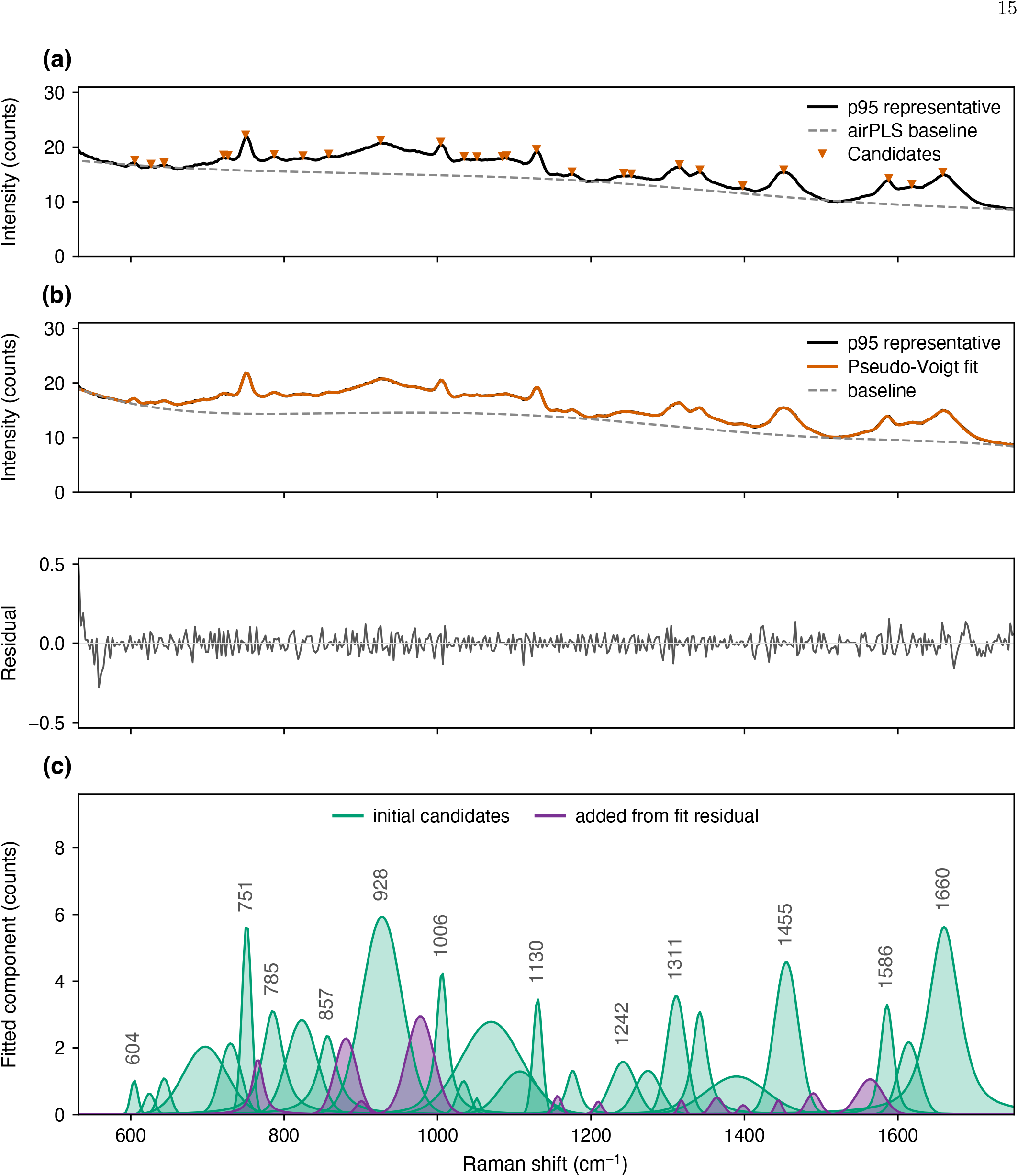
Data-driven dictionary construction (HeLa, 1 s). (a) 95th-percentile representative spectrum with airPLS initial baseline and detected candidates. (b) Bounded multi-peak pseudo-Voigt fit with locally readjusted baseline; residual below. (c) Final 38-band basis.

### B. Dictionary-constrained autoencoder

Each pixel spectrum is modeled as

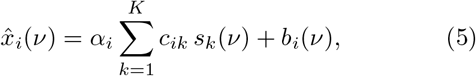

where the component spectra are linear combinations of the dictionary bands,

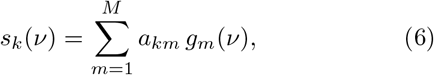

with simplex abundances *c*_*ik*_ *≥* 0, Σ_*k*_ *c*_*ik*_ = 1 (soft-max), a positive per-pixel gain *α*_*i*_ (softplus) that represents the total intensity, nonnegative dictionary amplitudes *a*_*km*_ (softplus) on the *fixed* basis *g*_*m*_ of Sec. III A, and a pixel-dependent smooth background *b*_*i*_ = **B***z*_*i*_ on a six-element cubic B-spline basis whose knot spacing (*~*400 cm^−1^) is well separated from the widest dictionary band (FWHM *≃*92 cm^−1^). A shared encoder (multilayer perceptron, MLP) predicts (*c*_*i*_, *α*_*i*_, *z*_*i*_) from the pixel spectrum; the component spectra *s*_*k*_ are global parameters. Endmembers are *l*_2_-normalized inside the decoder so that *α*_*i*_*c*_*ik*_ is the chemical amplitude. The training loss is the reconstruction term plus two weak regularizers, a quadratic penalty on the spline curvature and a (log *α*)^2^ gain prior. Two training modes are used. In least-squares training the encoder input and the reconstruction target are both the gain-corrected, SVD-denoised spectra, and the background coefficients *z*_*i*_ are unconstrained. In raw-domain quasi-likelihood training the encoder receives the gain-divided but *undenoised* count-equivalent spectra (normalized by their mean), whereas the decoder output is scored against the undivided count-equivalent observation through the objective of Eq. (2) with the gain fields in the forward model, 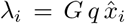; no SVD denoising enters this mode. Positivity of the rate is guaranteed by the parameterization: in this mode *z*_*i*_ is passed through a softplus so that *b*_*i*_ = **B***z*_*i*_*≥*0 on the nonnegative B-spline basis, *α*_*i*_ (softplus, floored at 10^−6^), *c*_*ik*_ (softmax), *a*_*km*_ (softplus), and *g*_*m*_ are all nonnegative, and *λ*_*i*_ is floored at 10^−8^ inside the logarithm, so that *λ*_*i*_ +*c >* 0 everywhere. No sparsity penalty is placed on the amplitudes *a*_*km*_: under the *l*_2_ normalization of the endmembers an *l*_1_ term on *a*_*k*_ would be scale-degenerate (a uniform rescaling of *a*_*k*_ leaves *s*_*k*_ unchanged). A vestigial *l*_1_ term of weight 10^−6^, inherited from the regular-grid variant in which it selects among redundant grid columns, was present in the runs reported here; it contributes less than 0.1% of the training loss, and repeating the 1 s decomposition without it leaves the fitted amplitudes (mean 0.83) and the cytochrome-*c* endmember correlation (0.92–0.93) unchanged, with map correlations identical to four decimals for two of three seeds and within run-to-run variation for the third (0.95 versus 0.98). Optional small bounded refinements of the dictionary (| Δ*µ*| *≤*3 cm^−1^, width scale 0.8–1.2) are implemented but off by default; they changed the results only marginally, confirming the accuracy of the fitted dictionary.

Two practical points matter for reproducibility. First, the softmax abundance head exhibits winner-take-all dynamics: components that lose all share (fraction of the total chemical amplitude) early in training become permanent “ghosts” that retain their initialization spectrum. Good initialization (NMF endmembers of a high-SNR dataset, transferred across exposure times and even cell lines) reliably avoids the degenerate optimum; with a poor initialization the same model collapses (e.g., shares 0*/*59*/*41 instead of 15*/*46*/*40 on C2C12). The initialization is always specified in spectral space, as *K* endmember spectra on the wavenumber axis; at model construction its smooth part is removed by projection onto the B-spline basis and the remainder is projected by nonnegative least squares onto the dictionary in use to obtain the initial amplitudes 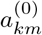. The same initialization can therefore be transferred to a dictionary of any size or origin (below: the 38-band HeLa and the 27-band C2C12 dictionaries). Second, the same mechanism produces an empirical capacity saturation: in a sweep of *K* = 2–10 on the 1 s data the number of components with nonzero share saturates at 3–6, reconstruction error is flat beyond *K* = 3, and component spectra remain stable (matched cosine 0.93–0.98 between adjacent *K*).

### C. Evaluation methodology

Beyond global map correlation, three metrics and one controlled test are used. (i) *Band-resolved fidelity* : a map is decomposed into difference-of-Gaussian bands,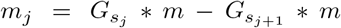, with Gaussian widths *s*_*j*_ = 0.1, 0.2, 0.4, 0.8, 1.6, 3.2 µm; for each band we report the Pearson correlation between the estimated and the reference band maps and the amplitude-retention ratio std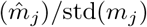, because a global correlation coefficient is amplitude-blind and rewards blur. (ii) *Crosstalk audit* : for each estimated component map the partial correlation with its own reference map is computed by regressing both maps, by ordinary least squares with an intercept, on the reference maps of the other components (and, on the phantom, on the true background amplitude) and correlating the residuals; this reveals leakage that diagonal correlations hide. The *spike fraction* of an endmember is obtained by detecting peaks in the unit-maximum-normalized spectrum with a prominence threshold of 0.05, measuring their widths at half prominence in cm^−1^, and counting the fraction narrower than 8 cm^−1^. (iii) *Time-matched self-consistency* : for living specimens, two statistically independent estimates from interleaved acquisitions are compared with each other rather than with a reference from a separate acquisition, at several lags; for learned denoisers the self-consistency must further be cross-checked against an independent estimator that does not share the learned inductive bias (a per-pixel or averaging estimator, even if noisier), because two networks with similar inductive bias agree with each other more than with the truth. (iv) *Known-truth phantom*: a noise-free intensity cube derived from the measured 1 s decomposition is passed through the calibrated detector model at several photon budgets, so that every estimator can be scored against truth and audited against the true background amplitude (Sec. IV F).

### D. AI-assisted analysis

Claude Code (Anthropic; model Claude Fable 5, `claude-fable-5`; used in August 2026) was employed to implement and audit the data-analysis scripts (preprocessing, gain-field estimation, dictionary construction, the dictionary-constrained autoencoder, the quasi-likelihood fits, the evaluation metrics, and the phantom simulations), to generate the figures, to retrieve and verify bibliographic metadata, and to draft and edit the manuscript text under the direction of the authors. The analysis objectives, the comparisons and parameter ranges, and the interpretation of all outputs were specified and independently reviewed by the authors; AI-generated code and numerical results were checked against the underlying data by the authors, who take full responsibility for the content.

## IV. RESULTS

### A. Decomposition of 1 s HeLa data

At *K* = 3 the model decomposes the (background-unsubtracted) HeLa cube into a cytochrome-*c* component with a clean resonance fingerprint and a mitochondrial network map, a broad 928 cm^−1^ “pedestal” component, and an amide-I/CH protein–lipid component [Fig. 3(a,b)]. The amplitude table *a*_*km*_ of each component reads directly as spectroscopy: for the cytochrome-*c* component the heme bands rank 751 *>* 1586 *>* 1311/1318 *>* 1130 cm^−1^ in area amplitude [Fig. 3(c)], accompanied by weak protein bands discussed below.

**FIG. 3.**
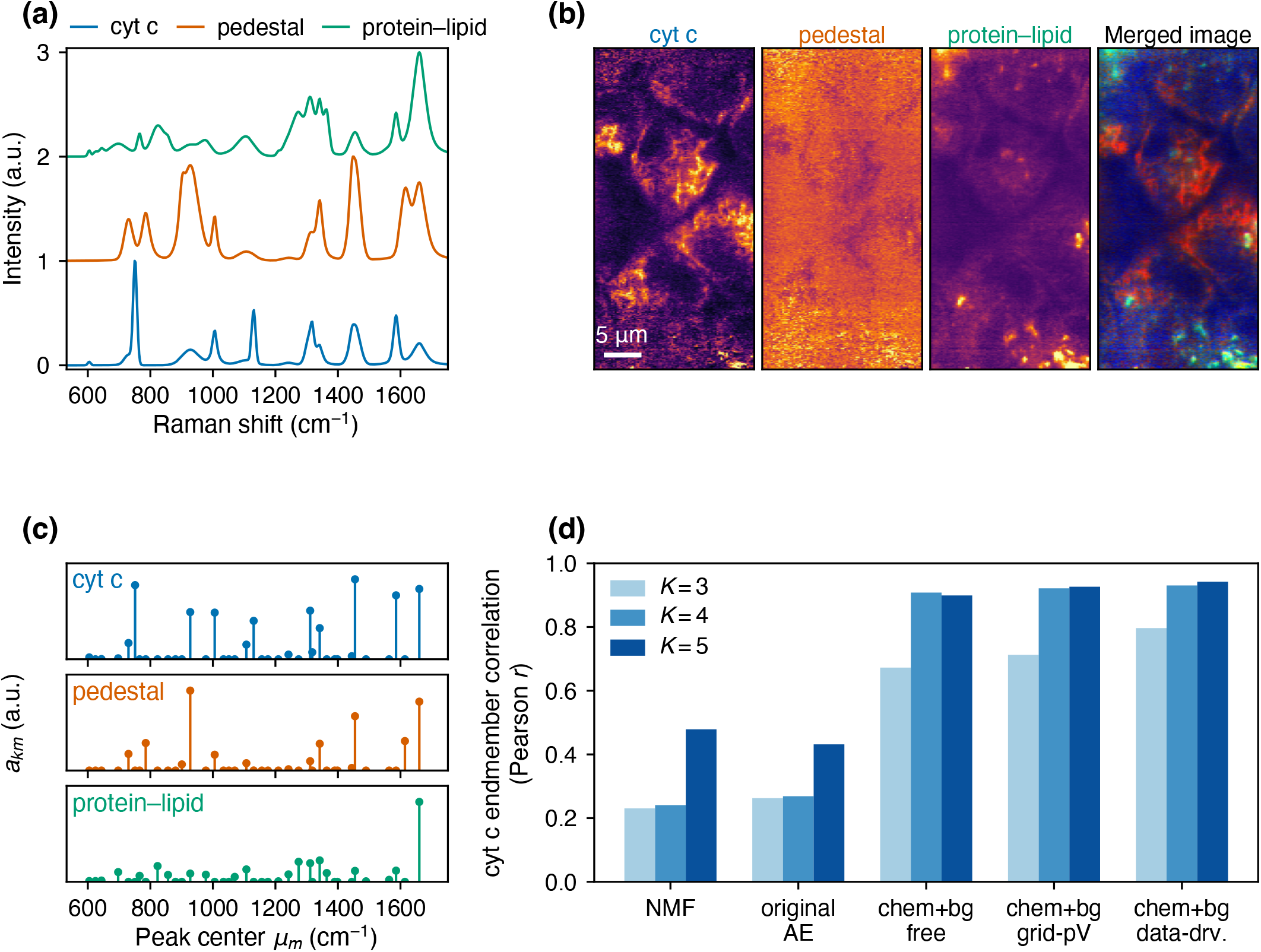
Dictionary-constrained decomposition of the 1 s HeLa cube (*K* = 3). (a) Component spectra (unit-maximum normalized and offset by one unit). (b) Abundance maps: cytochrome *c*, pedestal, protein–lipid, and their merged image (red: cytochrome *c*; green: protein–lipid; blue: pedestal). (c) Dictionary amplitudes *a*_*km*_ of the cytochrome-*c* component (stem plot). (d) Cytochrome-*c* endmember purity versus *K* for NMF, the unconstrained autoencoder, the free chemical+background decoder, the regular-grid dictionary, and the data-driven dictionary; spike fractions and *K*-stability values are given in the text.

*Assignment of the cytochrome-c component*. Reduced cytochrome *c* has Q-band absorption maxima at 550 nm (*α*, 0–0) and 520 nm (*β*)^18^; Q-band excitation enhances predominantly the Herzberg–Teller-active non-totally-symmetric heme modes—the anomalously polarized *A*_2*g*_ bands *ν*_19_, *ν*_21_, *ν*_22_ near 1585, 1313, and 1130 cm^−1^ and the depolarized *B*_1*g*_ pyrrole-breathing mode *ν*_15_ near 750 cm^−113,19,20^—while the oxidized form scatters roughly an order of magnitude more weakly under green excitation^3,21^. Our 532 nm line lies within the Q-band envelope, between the *β* (520 nm) and *α* (550 nm) maxima^18^, i.e. under the same resonance condition as the cellular studies at 532 nm^3,21,22^. Table I compares the component’s band amplitudes with literature spectra. Its four strong narrow bands, 751, 1130, 1311/1318, and 1586 cm^−1^, coincide within 3 cm^−1^ with the ferrocytochrome-*c* bands at 753, 1132, 1313, and 1585 cm^−1^ in solution at 514.5 nm^13^ and with the 750, 1127, 1314, and 1585 cm^−1^ bands observed in living cells at 532 nm^3,21,22^, the 750 cm^−1^ band being the strongest as in the cellular spectra. Bands that are strong in the 514.5 nm solution spectrum but absent here (*ν*_20_ (1400), the depolarized 1175 band, *ν*_11_ (1547), and *ν*_10_ (1622 cm^−1^); *ν*_4_ (1363) is at most a weak shoulder) are likewise absent or weak in the 532 nm cellular spectra of Refs.^3,21,22^ (only Ref.^22^ lists 1363), consistent with the excitation-profile dependence of the individual modes^13^ and with the lower signal-to-noise ratio of cellular spectra. The component is not a pure heme spectrum: it also contains weak, broad, non-resonant protein bands (phenylalanine 1006, CH deformation 1455, amide I 1660 cm^−1^; peak heights 0.2–0.4 of the 751 line, area amplitudes comparable to the heme bands because of their 3–4× larger widths) and a 1342 cm^−1^ band possibly from *b*-type cytochromes^23^. This is expected for mitochondria imaged at 250 nm pitch, where the resonant cytochrome-*c* signal is co-localized with the mitochondrial protein matrix; we therefore read the component as “cytochrome *c* within mitochondria” rather than as a molecular spectrum, with the filamentous perinuclear distribution of Fig. 3(b) as its spatial counterpart. A colocalization experiment with a mitochondrial stain would make the assignment definitive (Sec. V).

**Table I.** Bands of the cytochrome-*c* component compared with literature resonance Raman bands of reduced cytochrome *c* (730–1700 cm^−1^). Our bands are the dictionary centres *µ*_*m*_ with the NNLS amplitude *a*_*km*_ of the component on the area-normalized dictionary, scaled to max_*m*_ *a*_*km*_ = 1; bands below 5% of the maximum are listed only when the literature reports a band there (centre in parentheses). Literature positions: horse-heart ferrocytochrome *c* in solution, 514.5 nm (Ar^+^/Kr^+^ lines) with polarization ip (inverse/anomalously polarized, 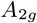) or dp (depolarized, 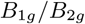) and intensity W/M/S/VS^13^; live cells or tissue at 532 nm^3,21–23^; Q-band-excited cytochrome *c* at 514.5 nm^24^; solution at 488 nm^25^. Mode labels follow the porphyrin numbering used in Refs.^19,20,24^. “—”: absent in that set.

| Our band (cm<sup>-1</sup>) | $a_{km}$ | Literature band (cm<sup>-1</sup>) | Assignment | Source(s) |
| --- | --- | --- | --- | --- |
| 730.0 (broad, FWHM 32) | 0.20 | — | not a heme band; low-frequency wing of the 751 line | — |
| 751.2 | 0.93 | 753 (dp, M); 750; 750; 750; 747 | $\nu_{15}$, $B_{1g}$ pyrrole breathing | 3,13,21,22,24 |
| 927.6 (broad, FWHM 61) | 0.59 | — | protein C–C backbone / cellular pedestal (non-resonant) | — |
| (977.5) | 0.00 | 975 (dp, W) | unlabelled dp heme band | 13 |
| 1005.8 | 0.58 | — | phenylalanine ring breathing (protein, non-resonant) | — |
| 1107.1 (broad, FWHM 59) | 0.18 | — | unassigned broad protein/lipid feature | — |
| 1130.5 | 0.48 | 1132 (ip, M); 1127; 1125; 1127; 1126 | $\nu_{22}$, $A_{2g}$ (third ip band; $C_\beta$–CH<sub>3</sub> vibration)<sup>a</sup> | 3,13,21–23 |
| (1156.5) | 0.00 | 1150 (dp, W) | unlabelled dp heme band | 13 |
| (1176.1) | 0.00 | 1175 (dp, S) | unlabelled dp heme band (strong at 514.5 nm) | 13 |
| 1241.9 (broad, FWHM 39) | 0.06 | 1228 (dp, M); 1248 | ours: amide III (protein); lit.: dp heme band | 13,22 |
| 1311.2 / 1317.9 | 0.61 / 0.09 | 1313 (ip, VS); 1314; 1310; 1310; 1313 | $\nu_{21}$, $A_{2g}$ (anomalously polarized) | 3,13,22–24 |
| 1341.7 | 0.39 | — (cyt. *c*); 1337 (*b*-type cytochromes); 1352 | not a ferrocytochrome-*c* band; *b*-type cytochrome and/or protein CH deformation | 22,23 |
| (1364.2) | 0.00 | 1365 (dp, M); 1363; 1363 | $\nu_4$ oxidation-state marker (ferrous 1363, ferric 1374; listed dp in Ref.<sup>13</sup>) | 13,22,25 |
| (1398.3) | 0.00 | 1400 (ip, S); 1398 | $\nu_{20}$, $A_{2g}$ (anomalously polarized) | 13,24 |
| 1454.6 (broad, FWHM 37) | 1.00 | — | CH<sub>2</sub>/CH<sub>3</sub> deformation (protein/lipid, non-resonant) | — |
| (1563.8) | 0.00 | 1547 (dp, M); 1548 | $\nu_{11}$ (dp) | 13,25 |
| 1585.9 | 0.80 | 1585 (ip, VS); 1585; 1585; 1582; 1584; 1582 | $\nu_{19}$, $A_{2g}$ (methine-bridge vibration; anomalously polarized) | 3,13,21–24 |
| — | — | 1600 (dp, W) | unlabelled dp heme band | 13 |
| (1614.2) | 0.00 | 1622 (dp, M); 1622; 1632 | $\nu_{10}$ (dp) | 13,22,25 |
| 1660.3 (broad, FWHM 47) | 0.88 | — | amide I (protein, non-resonant) | — |

<sup>a</sup> The $\nu_{22}$ label is inferred: Refs.<sup>13,20</sup> identify the strong Q-band bands as $\nu_{15}$ ( $B_{1g}$ ) and $\nu_{19}$ , $\nu_{21}$ , $\nu_{22}$ ( $A_{2g}$ ), and 1582/1313 are $\nu_{19}/\nu_{21}$ <sup>24</sup>; the $C_\beta$ –CH<sub>3</sub> description is from Ref.<sup>22</sup>.

The dictionary constraint is compared against the same architecture with (i) an unconstrained nonnegative decoder and (ii) a regular-grid pseudo-Voigt dictionary (centers every 6 cm^−1^, FWHM 6–120 cm^−1^) on identical data [Fig. 3(d)]. All three reach similar reconstruction error and map quality for cytochrome *c* (Pearson correlation *r* between abundance maps of 0.92–0.95 against a reference decomposition obtained independently of the autoencoder: MCR-ALS with *K* = 3 on the SVD-denoised, airPLS-background-subtracted 1 s cube, initialized by NMF; the same reference is used for the end-member purity of Fig. 3(d)), but they differ in spectral plausibility: the free decoder produces spiky endmembers (fraction of detected peaks narrower than 8 cm^−1^: 0.70– 0.82), the regular grid still contains grid-artifact structure, whereas the data-driven dictionary eliminates isolated spikes entirely (spike fraction 0.00; mean linewidth≃ 32 cm^−1^) and, unlike the alternatives, cannot place intensity at positions where the dataset has no band. It also gives the best stability against *K* (all-component matched cosine 0.935*/*0.976 for *K*=3→ 4→ 5, versus 0.79– 0.94 for the alternatives).

### B. Transfer to 100 ms-per-line imaging

The dictionary and initialization built at 1 s were applied unchanged to the 100 ms-per-line time-lapse (*~* 12 s per sweep; per-pixel signal *~*1.4 counts per channel). The result differs between spectra and maps: the cytochrome-*c* endmember is recovered with spectral correlation 0.92 to the 1 s result even from a *single* sweep, whereas the corresponding single-sweep map is photon limited (correlation 0.29 with the 1 s map at native resolution; the mitochondrial network becomes recognizable only after resolution-matched smoothing) [Fig. 4(a,b)]. Averaging the ten sweeps and applying a light resolution-matched smoothing (*σ* = 0.1–0.2 µm) yields map correlation 0.68 against the 1 s target while preserving the informative spatial bands, in contrast to strong smoothing of a single sweep, which raises the global correlation but destroys the 0.2–0.8 µm content (a metric artifact exposed by the band-resolved analysis of Sec. III C). Total-variation priors underperformed quadratic or Gaussian smoothing at every photon level tested; this behavior is consistent with edge information becoming unreliable at a peak SNR below 1 per pixel.

**FIG. 4.**
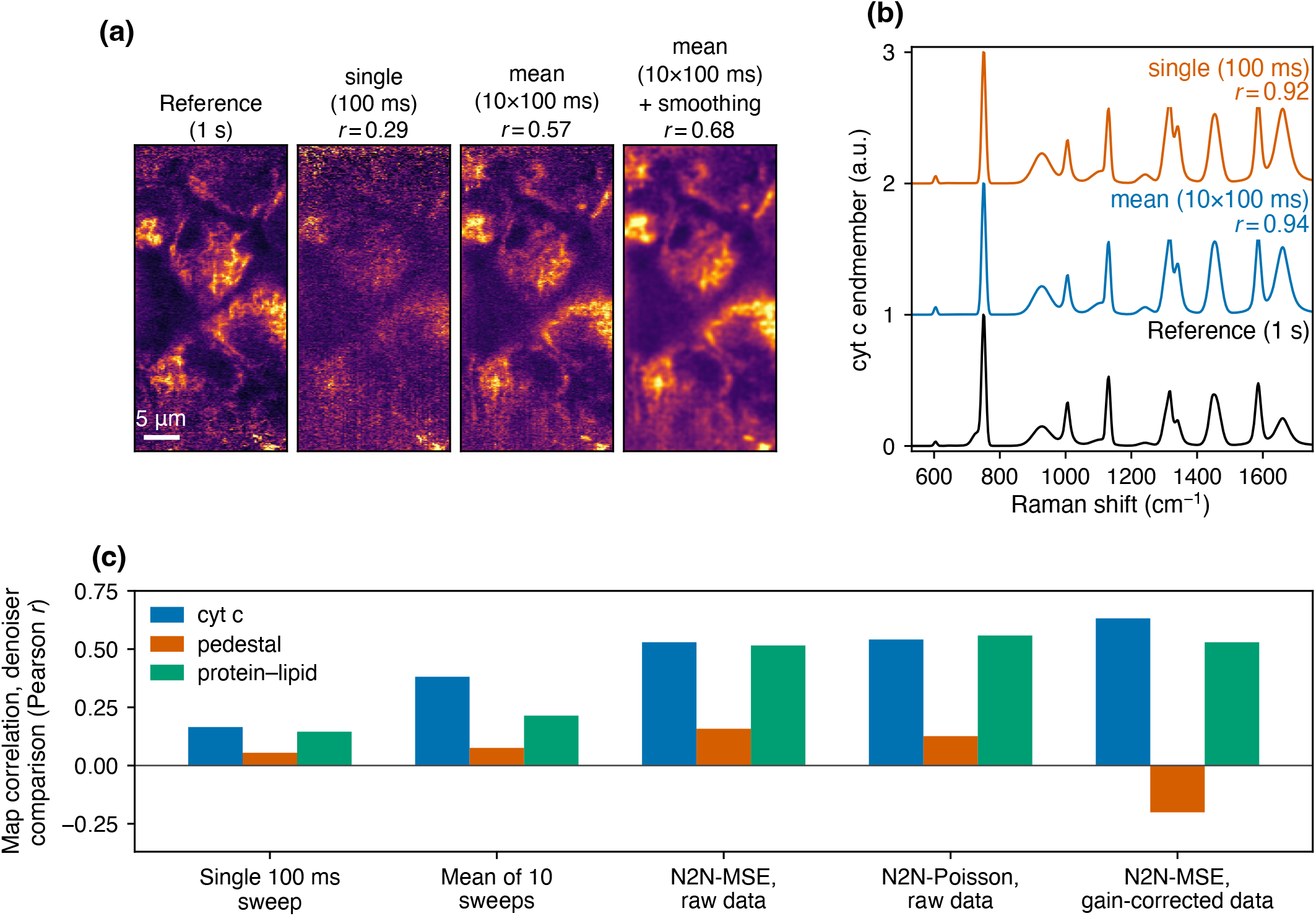
Transfer of the 1 s dictionary to 100 ms-per-line imaging. (a) Cytochrome-*c* maps: 1 s target; single 100 ms sweep; ten-sweep average; average with resolution-matched smoothing. (b) Cytochrome-*c* endmembers recovered at each photon level (unit-maximum normalized and offset by one unit). (c) Noise2Noise variants versus plain averaging (map fidelity, all three components), including training-domain and loss comparisons. Panel (c) is a self-contained comparison: all five variants, the two classical baselines included, were decomposed and scored under one protocol common to that comparison, which is not the per-condition pipeline of (a) and (b). The classical bars of (c) consequently differ in value from the *r* printed in (a); only differences within (c) should be read.

Noise2Noise denoising^14^ across sweeps is included as a case study of learned denoising rather than as part of the recommended pipeline. It raises the apparent single-sweep cytochrome-*c* map correlation with the 1 s target to 0.63 (network with spatial–spectral context input), against 0.38 for the ten-sweep average and 0.17 for the undenoised single sweep [Fig. 4(c)]. All bars of Fig. 4(c), the two classical baselines included, belong to this separate denoiser comparison, in which every variant is decomposed and scored under one protocol common to that comparison; they are internally comparable but systematically lower than, and not interchangeable with, the values in Fig. 4(a), which are obtained from the recommended pipeline trained separately at each photon level. The improvement does not carry over uniformly to the other components, and Sec. IV D shows that such agreement must itself be audited for shared bias. The choice of training domain matters more than the loss: networks trained on gain-corrected data give the best cytochrome-*c* maps but corrupt the broad components (partial correlations turn negative), whereas raw-domain training under the shifted-Poisson loss of Eq. (2) is the most spectrally faithful for multi-component quantification and produces calibrated rate estimates (held-out Pearson residual std 0.99). The gain of the shifted-Poisson quasi-likelihood over the mean-squared-error (MSE) loss in the raw domain is small but consistent (Δ*r ≃* 0.01–0.04).

The photon budget is strongly component dependent: the cytochrome-*c* component keeps its spectral identity from a single sweep and its map becomes usable with sweep averaging; the amide-I protein–lipid component needs sweep averaging plus smoothing; and the broad 928 cm^−1^ pedestal component, which is nearly degenerate with the smooth background, is not recoverable at 100 ms by any estimator tested (map *r ≤* 0.2).

### C. Contributions of the dictionary and of the reference initialization

The procedure of Sec. IV B transfers two distinct pieces of information from the 1 s data, the dictionary and the endmember initialization; their roles were separated by removing them in turn [Fig. 5]. The 100 ms analysis (ten-sweep average and single sweep; *K* = 3; three random seeds per condition; maps evaluated against the 1 s reference after resolution-matched smoothing, *σ* = 0.2 µm) was repeated at three levels of reference information: (A) the core dictionary plus the 1 s initialization, i.e. the recommended pipeline; (B) the dictionary alone, with the initialization derived from the 100 ms data themselves; and (C) neither, i.e. a free nonnegative decoder with self-derived initialization.

**FIG. 5.**
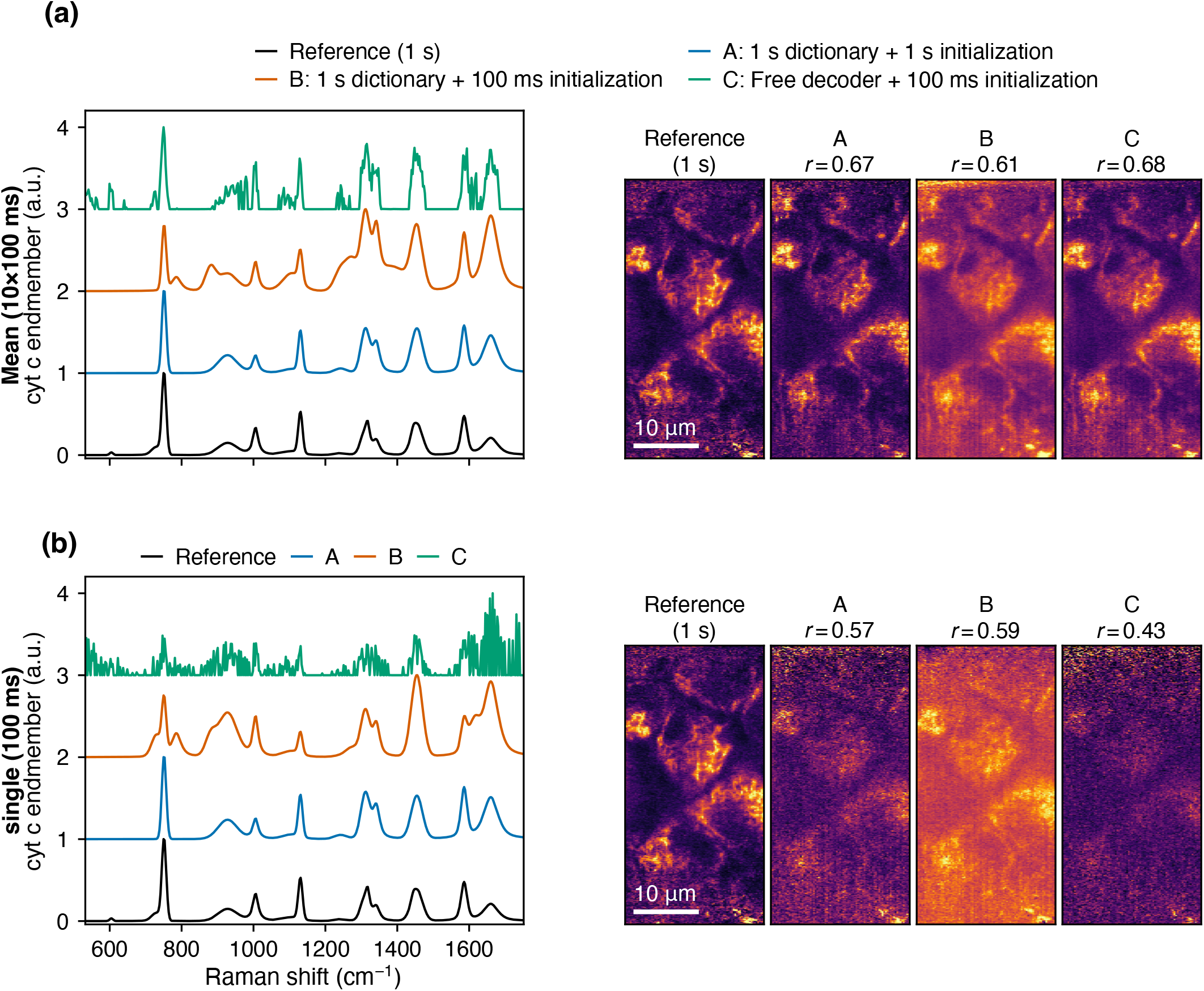
Effect of removing the transferred reference information on the 100 ms data (*K* = 3; median of three seeds). (a) Mean of ten 100 ms sweeps; (b) a single 100 ms sweep. Each row: cytochrome-*c* endmembers (unit-maximum normalized and offset vertically by one unit for clarity; 1 s reference and conditions A–C) and the corresponding maps with their correlation *r* to the 1 s reference (resolution-matched smoothing, *σ* = 0.2 µm). A: 1 s dictionary plus 1 s initialization (recommended pipeline); B: 1 s dictionary with 100 ms initialization; C: free nonnegative decoder with 100 ms initialization. Condition B in both rows, and condition C in the single-sweep row, collapsed to two active components; their seemingly competitive cytochrome-*c* map correlations are produced by a degenerate mixed component (see text).

The two ingredients fail in different ways, and the failures are complementary. *The reference initialization controls structural identifiability*: with a self-derived initialization the decomposition collapsed to two active components in every seed of condition B (both datasets) and of condition C on a single sweep—on the ten-sweep average of condition B the protein–lipid component dies outright (share 0%; its map slot anti-correlates with the reference at −0.46), and on a single sweep up to 99% of the chemical amplitude concentrates in one mixed component. The one self-initialized case that kept three active components, the free decoder on the ten-sweep average (shares 27*/*49*/*24), pays for it spectrally: its cytochrome-*c* endmember correlates with the 1 s spectrum at only 0.83, against 0.94 for the same data with the 1 s initialization. Every run with the 1 s initialization, by contrast, retained all three components and recovered the cytochrome-*c* endmember at 0.93–0.94 even from a single sweep. Fully *fixing* the basis (estimating only abundances) gives the same map fidelity as initialization transfer (0.66 versus 0.67 on the average, 0.55 versus 0.57 on a single sweep): transferring the initialization is sufficient, and fixing is not required. *The dictionary controls spectral physicality* : without it the endmembers develop narrow-spike structure (fraction of fitted peaks narrower than 8 cm^−1^: 0.67–0.94; median linewidth 3–5 cm^−1^), and the single-sweep “endmember” is dominated by noise (correlation 0.44 to the 1 s spectrum), even though the maps of the strong cytochrome-*c* component remain competitive on the averaged data.

This comparison also provides a concrete example of the evaluation pitfalls of Sec. III C: the degenerate two-component solutions still score global cytochrome-*c* map correlations of 0.43–0.62, the two dictionary-based ones (0.59 and 0.61) as high as the intact pipeline (0.57 and 0.67), because a dominant mixed component tracks the total cellular signal. The collapse is invisible to the global correlation and is exposed only by the share diagnostics and the partial-correlation crosstalk audit (the dead component’s partial correlation is zero or negative). A single fidelity number, reported without a structural audit, would have ranked a qualitatively broken decomposition equal to the working one.

### D. Photon limits versus acquisition mismatch

Increasing the number of averaged 100 ms sweeps improves the protein–lipid map only up to *n* ≃ 6 and then saturates at *r* ≃0.43 against the 1 s reference, far below the photon-scaling extrapolation [Fig. 6(a)], i.e. the correlation *r*(*n*) = (1 + *b/n*)^−1*/*2^ expected if the map error were pure photon noise averaging as 1*/n*, where *b* is the ratio of the single-sweep noise variance to the signal variance; *b* ≃ 13 is obtained by a least-squares fit of 1*/r*^2^− 1 = *b/n* to the classical track. The saturation is not caused by reference noise (the noise-limited self-consistency of the 1 s reference, from independent odd/even spectral-channel halves, is 0.98), by registration (shift scans are flat), or by the estimator. The cause can be identified by time-matched self-consistency. The protein–lipid maps of individual sweeps, audited by partial correlation against the other components, agree with each other at 0.85–0.91 for every lag from 12 to 109 s (trend *−*0.003 *±*0.003 per sweep) [Fig. 6(b)], and the means of the two half-datasets agree at 0.97: within the 122 s time-lapse the component is stable, and it is reproducible at *~*0.9 from the 0.1 s of photons per pixel contained in one sweep and at 0.97 from the 0.5 s contained in five. The saturation against the 1 s map therefore reflects a mismatch between the two acquisitions (separate rolling scans (Sec. II), a 6 min delay between them, and a slow global drift of the specimen described below) rather than the photon budget or a fast reorganization of the specimen. Whether cellular components reorganize on shorter time scales cannot be decided from these data; none is seen within 109 s.

**FIG. 6.**
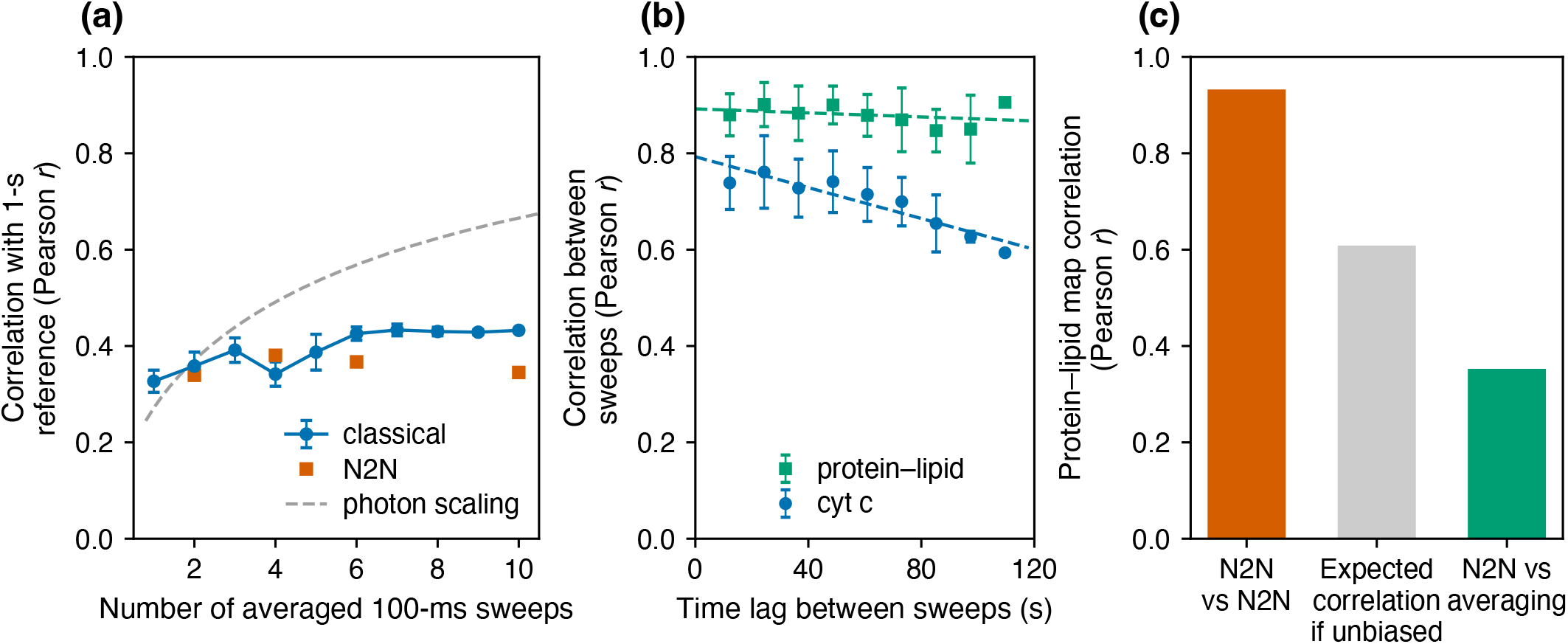
Photon limits versus acquisition mismatch. (a) Protein–lipid map fidelity versus number *n* of averaged 100 ms-per-line sweeps against the 1 s reference: photon scaling (dashed, (1 + *b/n*)^−1*/*2^ with *b* = 13 fitted to the classical track) followed by saturation. (b) Lag-resolved time-matched self-consistency (partial correlation against the other components) of the protein– lipid and cytochrome-*c* maps of individual sweeps: the protein–lipid component is stable over 109 s, so the saturation in (a) is acquisition mismatch, not photons or fast dynamics. (c) Noise2Noise self-consistency versus independent cross-checks: apparent agreement is inflated by shared bias.

Two consequences follow for the role of the 1 s data, which is itself a 102 s rolling acquisition recorded 6 min earlier (Sec. II). Its spectral products, the dictionary and the endmember initialization, are integrated over the field and comparatively insensitive to spatial rearrangement.

As a *map* reference, however, it is a separate acquisition: for the protein–lipid component, whose map differs between the two acquisitions, it cannot serve as a truth, and the correlations against it quoted in Secs. IV B and IV C are lower bounds set by acquisition mismatch rather than by the estimators. The cytochrome-*c* network drifts slowly but measurably: fixed-basis maps from the individual sweeps correlate with the 1 s map at 0.71–0.75 after resolution matching for the first 73 s of the time-lapse and at 0.57 at 109 s, the time-matched self-consistency of the cytochrome-*c* maps decays with a time constant of order minutes, and the drift is global (*~* 0.25 µm lateral and *~*20% in amplitude over 122 s), with the three column-thirds of the field varying in parallel so that no signature of the rolling reference acquisition is detectable. This slow drift, together with the 6 min inter-acquisition delay, is why the 1 s map is used as a reference only for cytochrome *c* and only with resolution matching.

The same time-matched comparison reveals a weakness of learned denoisers. Let *ρ*_self_ denote the agreement of an estimator with an independent copy of itself, obtained by applying it separately to the two half-datasets and correlating the two maps at native resolution, and *ρ*_cross_ the agreement between two *different* estimators. Two independently trained Noise2Noise networks are self-consistent at *ρ*_self_ = 0.93 (0.93–0.98 over the variants tested), apparently exceeding the corresponding figure for the 1 s reference itself (*ρ*_self_ = 0.875; the 0.98 quoted above is a different, purely noise-limited split of that reference, into odd and even spectral channels), whereas plain sweep averaging is self-consistent at only *ρ*_self_ = 0.40 on the same half-datasets. For two unbiased estimators with independent noise the cross-correlation equals the geometric mean of their self-consistencies, here 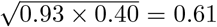; the measured network-versus-averaging value is only *ρ*_cross_ = 0.30–0.40 (mean 0.35), far below that [Fig. 6(c)]. The excess self-consistency is shared inductive bias, not information; the independently cross-checked native-resolution fidelity remains *~*0.6 for 0.5 s of photons regardless of the denoiser.

### E. Generality and minimal dictionaries

Applied to C2C12 myoblasts (four fields, 1 s), the identical pipeline (with a dictionary rebuilt from the C2C12 data in one minute (27 bands, representative-fit *R*^2^ = 0.998) and the HeLa endmember *spectra*, projected onto this 27-band dictionary as described in Sec. III B, as initialization) recovers the same three-component structure: the cytochrome-*c* endmember matches the HeLa one (cosine similarity 0.96), and per-field maps show mitochondrial networks including a pronounced perinuclear ring [Fig. 7(a,b)]. At *K* = 5 an additional, C2C12 protein-rich component emerges (27% share) dominated by phenylalanine 1003 and backbone C–C bands, consistent with myofibrillar protein. Wavenumber calibration transferred within half a channel (phenylalanine found at 1004.1, cytochrome *c* at 749.8 cm^−1^), and the HeLa and C2C12 dictionaries are interchangeable in practice: cross-applied, each reconstructs the other dataset’s representative spectra within 0.6–0.9% relative residual, with 19 of 27 bands shared.

**FIG. 7.**
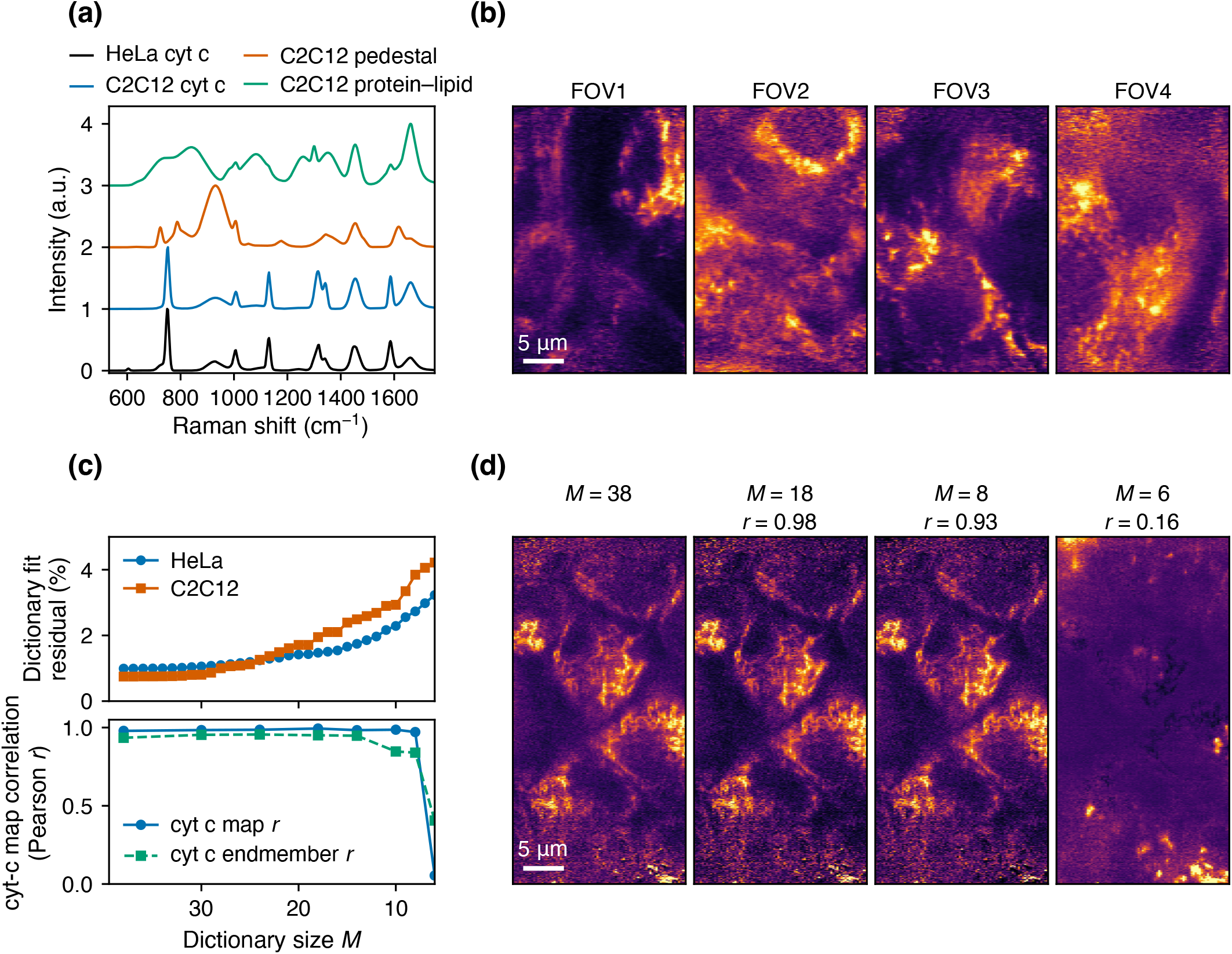
Generality and minimal dictionaries. (a) Unit-maximum-normalized C2C12 component spectra versus the HeLa cytochrome-*c* endmember (one-unit offsets). (b) C2C12 cytochrome-*c* maps (four fields). (c) Greedy dictionary pruning: representative-spectrum reconstruction and decomposition fidelity versus band count *M* ; the recommended *M* = 18 core and the collapse at *M* = 6 upon removal of the 751 cm^−1^ band. (d) Cytochrome-*c* maps at *M* = 38, 18, 8, and 6.

Greedy backward pruning quantifies how small the dictionary can be [Fig. 7(c,d)]. Decomposition quality, measured as the cytochrome-*c* map correlation with the standard *K* = 3 decomposition of Sec. IV A (trained independently), is flat down to *M* = 18 bands (0.988 ±0.005 versus 0.981± 0.003 for the full dictionary retrained within the same sweep, mean ±s.d. over three seeds; indistinguishable in map fidelity, with a small gain in end-member purity, 0.946± 0.005 versus 0.930± 0.005, from the removal of mild overcompleteness), degrades gently to *M* = 8, and collapses abruptly at *M* = 6, at the step at which the 751 cm^−1^ resonance band is removed: the cytochrome-*c* structure then migrates into the wrong component in both cell lines and at both exposures. A single-band deletion control separates the role of the 751 cm^−1^ band from the cumulative loss. Removing 751 alone from the 18-band core dictionary, with the 1 s initialization and three seeds, lowers the cytochrome-*c* endmember correlation from 0.95 to 0.72 (full dictionary: from 0.93 to 0.74) while the map correlation (0.96; full dictionary 0.95) and the three-component structure are retained; removing any other single cytochrome-*c* band (1130, 1311/1318, 1586 cm^−1^) or the phenylalanine band costs only 0.03–0.07, and removing the CH deformation band (1455 cm^−1^) costs 0.16 while shifting 34% of the share into the cytochrome-*c* slot. The 751 cm^−1^ band is thus the single most influential band for the spectral identity of the component, but the collapse at *M* = 6 is cumulative rather than caused by its removal alone. At 100 ms the pruning tolerance is slightly narrower (map *r* declines monotonically by 0.04 from *M* = 38 to *M* = 8): at low photon numbers even minor bands carry useful discriminating counts. An eighteen-band core dictionary is therefore recommended across the tested exposure conditions. Applied unchanged to the C2C12 data (four fields, three seeds), the HeLa-derived core retains the cytochrome-*c* map and endmember of the C2C12 standard at 0.977± 0.003 and 0.983 ±0.001 (full HeLa dictionary: 0.990 and 0.984; the C2C12 dictionary itself: 0.994 and 1.000) with all three components active, but the protein–lipid endmember falls to 0.69 (0.83 with the full HeLa dictionary): the core transfers across cell lines for the resonance component, whereas a dataset-specific dictionary remains preferable for the non-resonant components. In all cases resonance fingerprints should be protected from pruning regardless of their fitted area.

### F. Quantitative evaluation on a phantom with known truth

All results so far are judged against decompositions of the same measured data. To score the estimators against a known truth we built a phantom from the measurement itself: the *K* = 3 decomposition of the 1 s HeLa cube (abundance maps and endmembers of Sec. IV A plus the per-pixel airPLS background, lightly smoothed to remove inherited shot noise) defines a noise-free intensity cube, which is passed through the detector model of Sec. II (gain fields and resampling from the working second-order shifted-Poisson noise model, *y* = Poisson(*λ*+*c*) *−c*) at photon budgets equivalent to 1 s, 100 ms, and 10 ms exposure, plus the mean of ten independent 100 ms realizations, with three noise realizations per budget. The phantom reproduces the real 1 s data to 1.8% (mean spectrum) and 3.2% (per-channel standard deviation) RMS. All phantom estimators operate on the same representation as the corresponding real-data analyses of Secs. IV A–IV C (known gains divided out, SVD denoising, *K* = 3, i.e. the least-squares-domain pipeline); the raw-domain quasi-likelihood fits with the gains in the forward model were not part of the phantom comparison. Dictionaries are rebuilt from the noisy 1 s-budget realization (35–37 bands), and the “calibrated” dictionary autoencoder takes its initialization from that realization, as in the real workflow. Endmember fidelity is quantified by the *spectral angle* between the estimated and the true endmember, that is, the angle between the two spectra regarded as vectors, i.e. the arccosine of their cosine similarity (0°: identical shape; 17° corresponds to a cosine similarity of 0.96, 57° to 0.54). Map fidelity is quantified by the Pearson correlation *r* with the true map after resolution-matched smoothing and by the normalized root-mean-square error (NRMSE, the RMS error divided by the RMS of the true map after a least-squares scale fit). Because the truth is known, each map can also be audited by partial correlation against the other truth maps *and the truth background amplitude* (*r*_aud_), and compared with a reference estimator that uses the true endmembers (per-pixel nonnegative least squares, NNLS). Figure 8 summarizes the results.

**FIG. 8.**
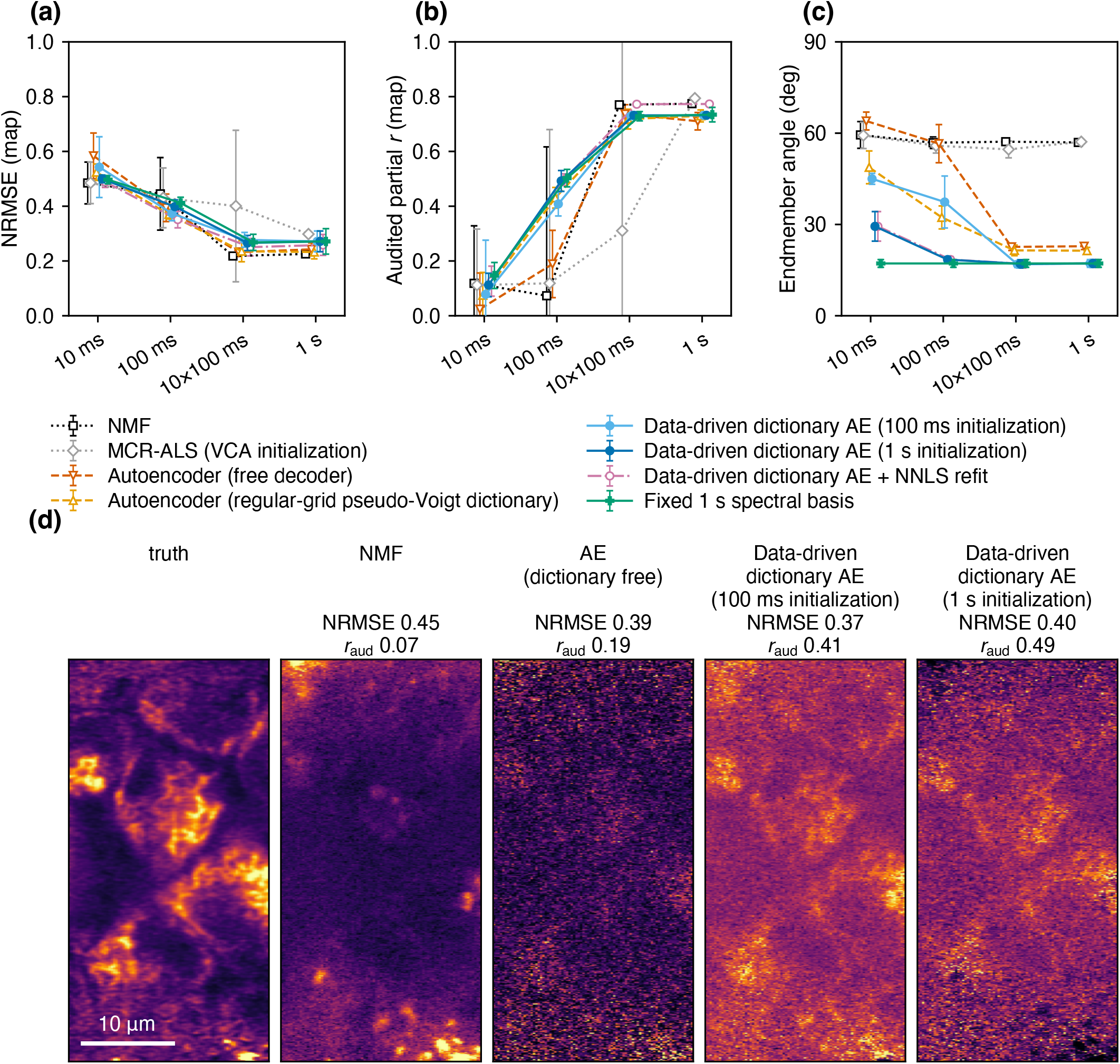
Measurement-derived phantom with known truth (three noise realizations; mean ±s.d.). (a) Cytochrome-*c* map NRMSE and (b) background-audited partial correlation *r*_aud_ versus photon budget for NMF, MCR-ALS, the dictionary-free and regular-grid autoencoders, and the dictionary-constrained autoencoder with 100 ms or 1 s initialization, with fixed basis, and with a final per-pixel NNLS refit. (c) Cytochrome-*c* endmember spectral angle. (d) Cytochrome-*c* maps at the 100 ms budget.

**FIG. 9.**
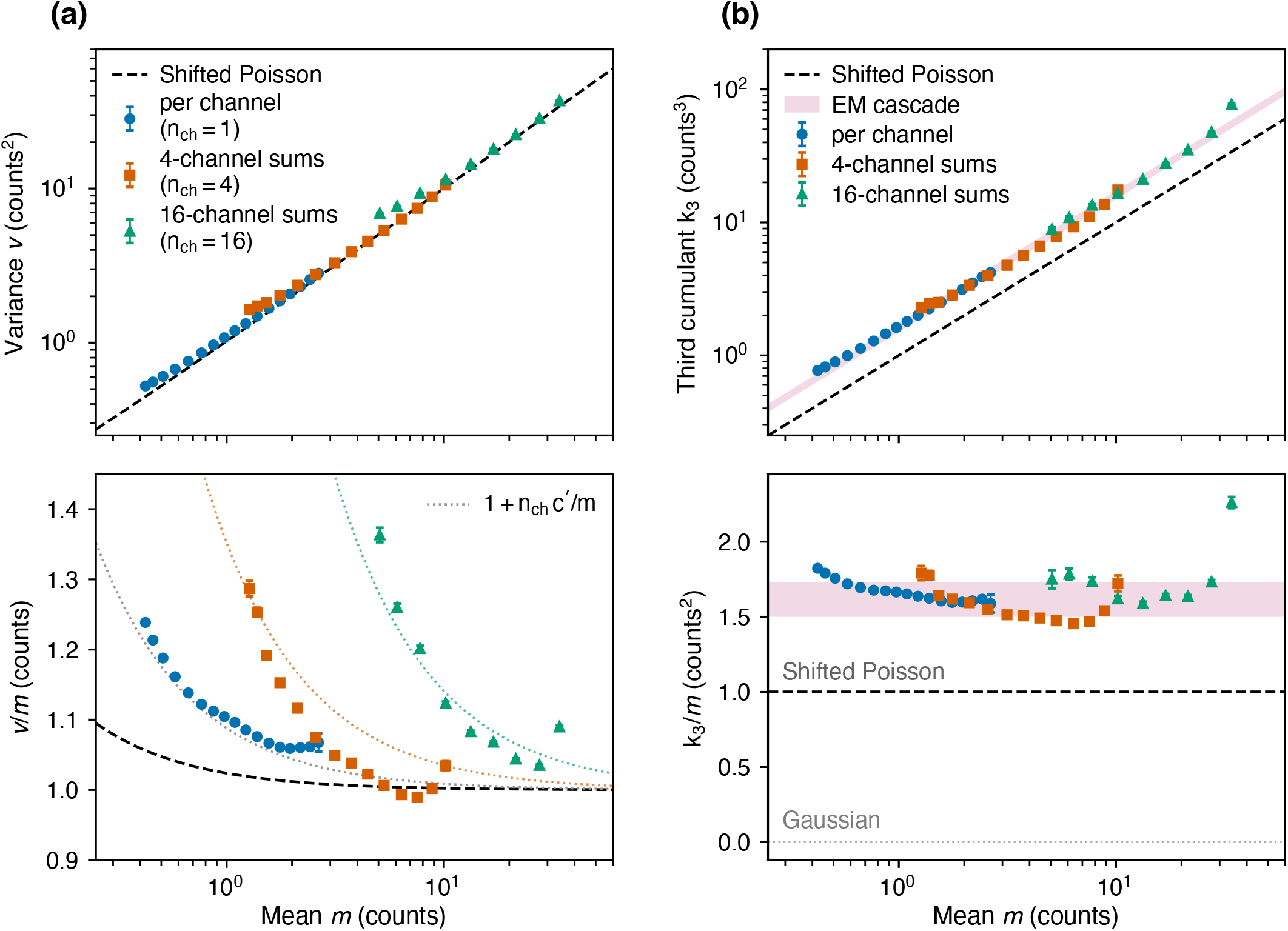
Detector-model diagnostics on the ten repeated 100 ms sweeps (triples binned on an independent intensity estimate). (a) Temporal variance *v* versus mean *m* per channel and for 4- and 16-channel sums, with the shifted-Poisson line *v* = *m* + *c*; lower panel, the ratio *v/m* showing the read-noise floor below*~* 1 count. The dotted curves are 1 + *n*_ch_*c*^*′*^*/m*, where *n*_ch_ is the number of summed adjacent channels and *c*^*′*^ = 0.088 counts^2^ is the fitted per-channel variance floor. (b) Third cumulant *k*_3_ versus *m* with the shifted-Poisson line (*k*_3_= *m*) and the electron-multiplication cascade band (*k*_3_= (6*/F*^4^) *m* for *F*^2^ = 2–1.86); lower panel, the ratio *k*_3_/*m*.

At the 1 s budget all estimators reach cytochrome-*c* map correlations of 0.85–0.95 and protein–lipid correlations of 0.90–0.96 (resolution matched), and the mean of ten 100 ms realizations is equivalent to a single 1 s realization for every method, as assumed in Sec. IV B. The highest raw correlation (NMF, 0.95) belongs, however, to a component whose spectrum is 57° from the true cytochrome-*c* spectrum: a mixed factor that tracks the total cellular signal. Audited against the background amplitude, all methods are at 0.71–0.79 at 1 s, and at the 100 ms budget the ranking inverts: NMF and MCR-ALS fall to *r*_aud_ = 0.07 and 0.12 (single realizations between− 0.55 and 0.48) and the free decoder to 0.19, whereas the dictionary estimators retain 0.41–0.50 with realization-to-realization scatter of only 0.04–0.06, at the level of the true-endmember reference (*r* = 0.65 versus 0.63– 0.76 raw). The classical “cytochrome-*c* maps” at 100 ms are thus total-intensity proxies, the same failure that the crosstalk audit revealed on real data in Sec. IV C.

The main contribution of the dictionary is endmember fidelity, at every budget of this measurement-derived phantom (whose truth lies within the model class; see Sec. V). The cytochrome-*c* spectral angle is 17–19° for the dictionary estimators down to≤ 100 ms (29° at ≤10 ms), against 57–64° for the free decoder at ≤100 ms (23° at ≥1 s equivalent), 32°/49° for the regular grid, and 55– 60° for NMF and MCR-ALS at every budget; the spike fraction is zero for every dictionary run against 0.55– 0.93 (free), 0.16–0.46 (grid), and 0.82 (NMF at ≤100 ms). Without the 1 s initialization the dictionary autoencoder loses spectral identity at low budgets (37 *±*9° at 100 ms and 45° at 10 ms, versus 18.5 and 29° with it), reproducing the spectral half of the real-data comparison of Sec. IV C; its structural half, however, is not reproduced: on the phantom every method keeps three active components at every budget. The two-component collapse observed on real data therefore reflects structure beyond a three-component-plus-smooth-background model (organelle motion between sweeps, background not spanned by the spline basis, minor species), not photon statistics. The component-resolved photon limits agree with Sec. IV B: cytochrome *c* is recoverable at 100 ms (*r*_aud_ ≃0.5) but not at 10 ms (≤ 0.15 for every estimator, including the true-endmember reference), the protein–lipid maps remain at 0.78–0.80 at 10 ms for the dictionary estimators (free decoder 0.62 ±0.23), and the broad pedestal is the weakest dictionary component at 100 ms, its broad bands being degenerate with the spline background. Fixing the basis is again not better than transferring the initialization (0.58 versus 0.63 raw, 0.50 versus 0.49 audited). One practical improvement was found: re-estimating the abundances per pixel by NNLS with the autoencoder’s own endmembers and background basis closes the gap between the amortized encoder estimate and a per-pixel refit: the 100 ms cytochrome-*c* map *r* rises from 0.63 to 0.76 and the NRMSE falls from 0.40 to 0.35 with the spectra unchanged. This refit is recommended as the final step of the pipeline.

## V. DISCUSSION

Three design rules follow. First, *measure what is measurable*: band centers, widths, and shapes are spectroscopic observables, and fixing them converts the ill-posed unmixing problem into amplitude estimation on a physically meaningful basis. The cost is a one-minute fit per dataset; the benefit is spike-free, directly interpretable endmembers, stability against the component number, and transferability. The comparison of Sec. IV C shows that the dictionary and the reference initialization have separate roles (spectral physicality and structural identifiability, respectively) and that neither failure mode is detected by global map metrics, so neither ingredient can substitute for the other. Second, *put known physics in the forward model and unknown noise in the quasi-likelihood* : for quasi-likelihood abundance estimation on the undivided count-equivalent output, known gain fields should remain in the forward model rather than being divided out of the measured counts, and denoising should not precede the fit; dictionary construction, by contrast, is an auxiliary measurement of band positions and widths for which gain correction and SVD denoising are appropriate. Keeping calibrated gains in the forward model and using the moment-calibrated shifted-Poisson quasi-likelihood yields calibrated residuals and the best multi-component consistency among the tested raw-domain losses, even though its point-estimate advantage over least squares is small (Δ*r* ≃0.01–0.04). Third, *evaluate against the right null* : for living specimens a reference from a separate acquisition drifts away, so that frame-sweep saturation against it separates the photon limit from acquisition mismatch while lag-resolved time-matched self-consistency shows what is recoverable; and self-agreement of learned estimators must never be mistaken for fidelity.

Limitations and outlook. All conclusions rest on single fields per condition of two adherent cell lines. The cytochrome-*c* assignment rests on the resonance band pattern, its agreement with literature spectra (Table I), and the filamentous perinuclear distribution (Sec. IV A); a colocalization experiment with a mitochondrial stain would make it definitive and is the obvious next step. The phantom of Sec. IV F quantifies the estimators against known truth, but its truth is generated within the model class (three components plus a smooth background): it reproduces the loss of spectral identity without the reference initialization but not the two-component collapse seen on real data, which therefore signals structure beyond that model, and the comparison is by construction favorable to an estimator whose dictionary spans the truth. The acquisition window excludes the silent region, where alkyne-tag labels^26^ and related nanoparticle Raman tags would add sharp exogenous fingerprints; a preliminary analysis of tagged samples in this study identified four distinct localized signatures, suggesting the dictionary approach extends naturally by appending measured tag bands. An exact detector likelihood (Poisson–Gaussian^15^ or the EM cascade^4^) is implemented as an interface, but the second-order shifted-Poisson quasi-likelihood was sufficient down to~ 1 count per channel. The softmax ghost-component instability, although manageable by initialization transfer, deserves a principled fix (e.g., annealed or over-parameterized abundance heads). Finally, the proposed time-matched evaluation suggests an acquisition design: interleaving short-exposure sweeps with sparse high-SNR reference sweeps would let the same experiment calibrate the dictionary, train the denoiser, and validate the maps.

## VI. CONCLUSIONS

A data-driven spectroscopic dictionary, frozen inside a chemical-plus-background autoencoder with detector-calibrated quasi-likelihoods, turns photon-limited Raman unmixing into interpretable band-amplitude imaging: calibrate once at high SNR, then image at 100 ms exposure per line. On live HeLa cells the approach eliminates artificial spectral structure, stabilizes the decomposition, and transfers across exposure time; its construction procedure together with the transferred initialization generalizes to C2C12 myoblasts; and it reduces to an eighteen-band core dictionary in which the 751 cm^−1^ resonance band is the most influential single element (its sole removal costs 0.2 in endmember purity, three times more than any other cytochrome-*c* band), although the collapse of the decomposition requires cumulative pruning. Equally important, we show that in live-cell, photon-limited imaging the evaluation methodology is part of the method: lag-resolved time-matched self-consistency and independent-estimator cross-checks change the conclusions about what is and is not recoverable.

## ACKNOWLEDGMENTS

This research was supported by JST FOREST (JP-MJFR216R), JSPS KAKENHI Grant-in-Aid for Scientific Research (B) (JP23K23297, JP25K03136), Grant-in-Aid for Transformative Research Areas (JP25H01396, JP25H01393), Grant-in-Aid for Challenging Research (Pioneering) (JP25K21710), Grant-in-Aid for Scientific Research (S) (JP25H00410), Grant-in-Aid for Scientific Research (A) (JP26H02285), the Photographic Research Fund of the Konica Minolta Imaging Science Foundation, the Murata Science and Education Foundation, the Research Foundation for Opto-Science and Technology, and the Inamori Research Grants Program of the Inamori Foundation. The use of an AI assistant in the data analysis and in the preparation of the manuscript is disclosed in Sec. III D.

## AUTHOR DECLARATIONS

## Ethics approval

Ethics approval is not applicable; the study used established cell lines only.

## Author contributions

**Shunsuke Yagi:** Formal analysis; Methodology; Software; Validation; Visualization; Writing – original draft; Writing – review & editing. **Norihide Sagami:** Formal analysis; Methodology; Software; Validation; Visualization; Writing – original draft; Writing – review & editing. **Ikuto Eshima:** Investigation; Data curation; Writing – review & editing. **Kotaro Hiramatsu:** Conceptualization; Supervision; Project administration; Funding acquisition; Formal analysis; Methodology; Software; Validation; Writing – original draft; Writing – review & editing.

## DATA AVAILABILITY

The analysis code (dictionary construction, dictionary-constrained autoencoder, evaluation protocols) and the raw and processed data supporting the findings of this study are available from the corresponding author upon reasonable request.

## Appendix A Model and training hyperparameters

Table II lists every setting of the dictionary construction and of the dictionary-constrained autoencoder used for the results in the main text; the same settings were used for all datasets and photon levels unless stated otherwise. Two settings were varied on the 1 s data to check sensitivity (single seed). The SVD denoising rank (4 adaptive, 6, 8, none) leaves the endmembers unchanged (cytochrome-*c* endmember correlation 0.935–0.939) while the native-resolution map correlation with the standard decomposition falls with retained noise (0.98, 0.94, 0.94,

**TABLE II.**
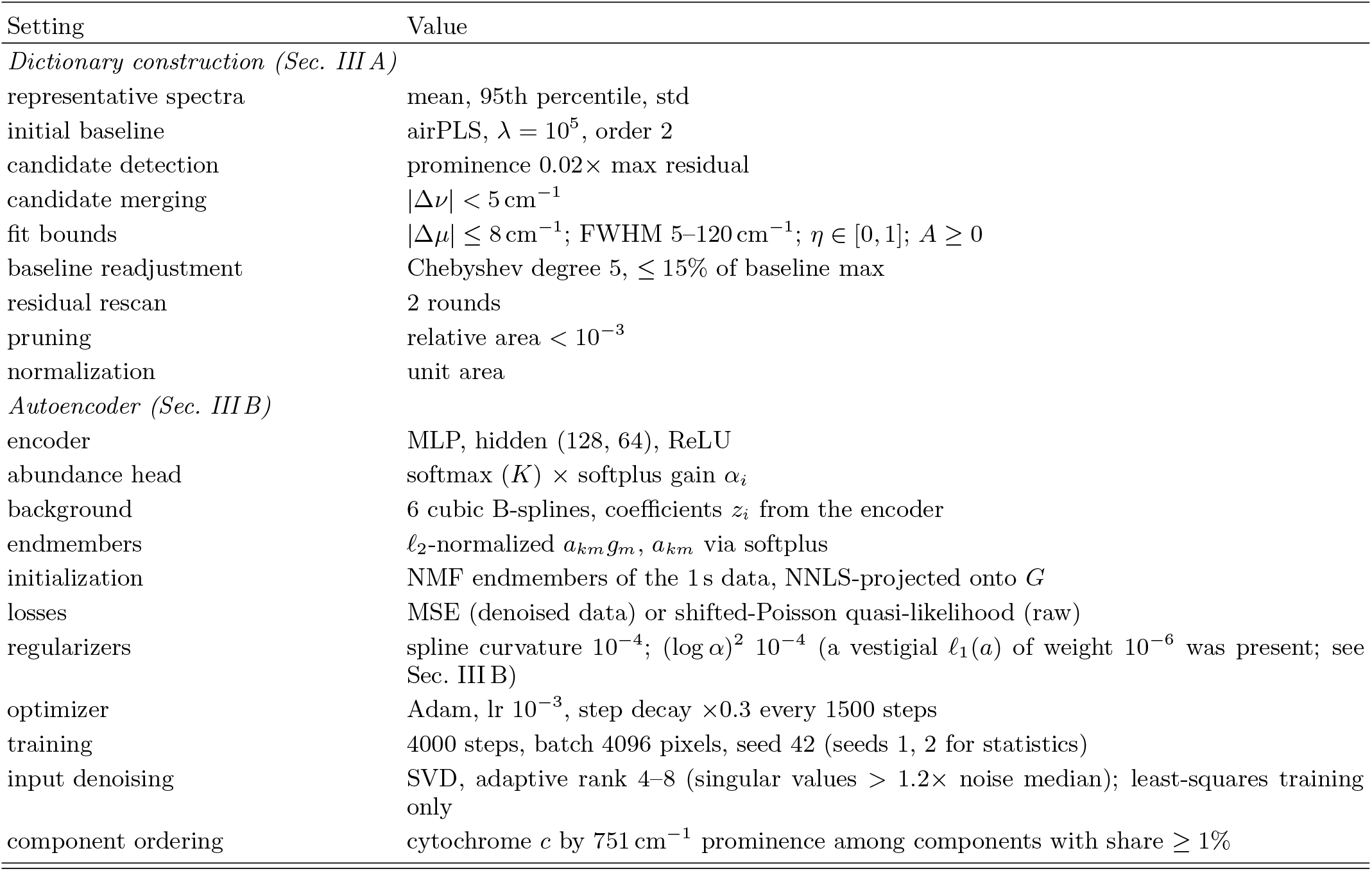
Dictionary-construction and autoencoder settings.

0.73). The peak-detection prominence (0.01, 0.02 default, 0.04 of the maximum residual) yields dictionaries of *M* = 50, 38, and 28 bands with cytochrome-*c* map correlations 0.98–0.99 and endmember correlations 0.91– 0.95; component shares shift by up to 7 percentage points at *M* = 28.

## Appendix B Detector-model diagnostics

Figure 9 shows the second- and third-moment diagnostics of the count-equivalent output that underlie Sec. II: the temporal variance and the third cumulant of the ten repeated 100 ms sweeps as functions of the mean, per channel and for sums of 4 and 16 adjacent channels (cumulants add, so the sums extend the intensity range to ~40 counts while preserving the slopes). The variance follows *m* + *c* with unit slope; the third cumulant follows the electron-multiplication cascade band (6*/F*^4^) *m* rather than the Poisson line *m*. The floor of the variance is 0.09 counts^2^.

